# DMS-MapSeq Analysis of Antisense Oligonucleotide Binding to lncRNA PANDA

**DOI:** 10.1101/2023.10.22.563486

**Authors:** Gabriel Romero-Agosto, Ethan Cox, Silvi Rouskin

**Affiliations:** Department of Microbiology, Harvard Medical School; Utah Valley University

## Abstract

While various methods exist for examining and visualizing the structure of RNA molecules, dimethyl sulfate-mutational profiling and sequencing (DMS-MaPseq) stands out for its simplicity and versatility. This technique has proven effective for studying RNA structures both in vitro and in complex biological settings. We’ve updated the protocol for using DMS-MaPseq, and it can also be employed to identify the binding of antisense oligonucleotides (ASOs) to RNA. By applying this updated protocol, we successfully characterized the structural ensemble of the HIV1 Rev Response Element (RRE), along with its two alternative structures. The findings align with previously published research. Additionally, we resolved the structure of the long non-coding RNA PANDA, which was previously unknown. Moreover, we used PANDA as a basis for designing ASOs and confirmed their binding through a substantial decrease in DMS-reactivities at the anticipated ASO binding locations.

## Introduction

It is abundantly clear that RNAs, particularly non-coding RNAs, play critical roles in regulating cellular processes. This functionality is possible thanks to the RNA’s ability to adopt secondary and tertiary structures^1^. These structure ensembles are relevant for the function of 3’UTRs of mRNA, microRNAs, lncRNA, and viral RNA^2-5^. Thus, being able to determine RNA structure is critical for understanding cellular regulatory mechanisms. Low-throughput methods such as X-ray crystallography, NMR, and Cryo-EM that have been used to interrogate RNA and its conformations^6^. Additionally, there are in-silico approaches that make use of thermodynamic-based algorithms for RNA structure prediction^7^. However, these methods are constrained to unphysiological environments (high salt, high RNA concentrations, low temperature etc.) and fail to consider factors such as RNA-binding proteins (RBPs), RNA modifications, crowding, and the specifics of the cellular environment.

Few techniques have been developed to assay RNA structures in biologically relevant environments like cells or tissues; chemical probing being the most renowned and attractive one. Chemical probing directly measures the base-pairing state of nucleotides. The two most famous methods are RNA structure probing with selective 2′-hydroxyl acylation analyzed by primer extension (SHAPE) and Dimethyl sulfate (DMS)^4^. DMS is perhaps the most commonly used RNA structure probing chemical, given that it selectively methylates unpaired adenines and cytosines within their Watson-Crick face^8-10^. In early days, DMS treatment was coupled with Reverse-transcription (RT)-PCR. The RT enzyme drops-off upon reaching modified bases, generating truncated cDNAs that could be visualized using gel electrophoresis, a technique known as DMS-foot printing^11,12^. However DMS footprinting evolved into DMS mutational profiling (MaP) with the development of highly processive RT enzymes^13,14^. Enzymes such as thermostable group II intron reverse transcriptase (TGIRT-III) read through the DMS modification and incorporate a mismatched base, which leads to a mutation in the cDNA sequence^10^. DMS-MaP enabled detection of multiple unpaired bases on a single RNA molecule. Improvements in high-throughput sequencing (HTS) coupled with DMS-MaP made it possible to determine the RNA structures of whole transcriptomes^15,16^.

DMS-MaPseq has been used in-tandem with Detection of RNA folding Ensembles using Expectation Maximization’ (DREEM) algorithm to identify alternative structures that form from the same RNA sequence^17^. Together, DMS-MaPseq and DREEM have been used to study conformation ensembles within the HIV-1 and SARS-CoV-2 genomes^10,17^. These together with other ensemble approaches such as DRACO^18^ and DANCE-MaP^19^ have been pivotal to the advancement of the RNA structure field; however, experimental advancements have made the previous DMS-MaPseq protocol outdated.

One of the main modifications made to the protocol is the change from TGIRT-III to NEB’s Induro^®^ RT, a commercially available group II intron-encoded reverse transcriptase. We validate this change using the HIV-1 Rev Response Element (RRE), a small RNA sequence that is recognized by the viral Rev protein for exportation of splice and unspliced HIV-1 transcripts.

DMS-MaPseq previously revealed that RRE adopts a five-stem and four-stem alternative structures in vitro and in physiological environments^17^. Since the RRE is a highly structured and well-studied sequence it serves as a great model to validate updates to the DMS-MapSeq protocol.

We also demonstrate DMS-MapSeq’s ability to identify the binding of antisense oligonucleotides (ASOs) to target RNA sequences. ASOs are a class of RNA targeting molecules that have recently gained attraction for their therapeutic potential. They have many mechanisms of actions to modulate protein levels^20^. These oligonucleotides can also undergo an array of modifications that fine tune mechanisms of action, affinity, durability, and modes of delivery^21^. To this day as many as ten genetic disorders have FDA approved ASO-based treatments^22^. Here we use DNA ASOs to target P21-Associated ncRNA DNA Damage-Activated (PANDA). PANDA is a lncRNA about 1,500 nucleotides long that is transcribed upon DNA-damage in a p53-dependat manner^23^. Like many lncRNAs, PANDA is capable of regulating cell-growth. It specifically interacts with cell factors to downregulate pro-apoptotic genes, favoring cell-cycle arrest^23,24^. Given its importance as a regulator for the DNA damage response, it comes to no surprise that its deregulation has been linked to cancer progression^25-27^. However, little is known as to how PANDA interacts with its binding partners and solving its structure will aid to better understand these interactions.

## Method Overview

The primary objective is to provide the reader with a detailed method on how to perform DMS-MaPseq on in-vitro transcribed. The steps outlined here are broadly applicable to most in-vitro generated RNA, but modifications will have to be made depending on the length of the molecule and its sequence. As a secondary objective, details as to how to disrupt RNA structures or test ASO targeting are provided as well. An important note is that many of these reagents can be substituted for other commercially available products. The essential reagents are stated explicitly.

## Method I: DMS-MaPseq on HIV1-RRE

### PCR of the HIV1-RRE

The HIV1_RRE was ordered as a 232-nucleotide segment in a gene-block from IDT. Primers were designed using SnapGene and ordered through IDT:

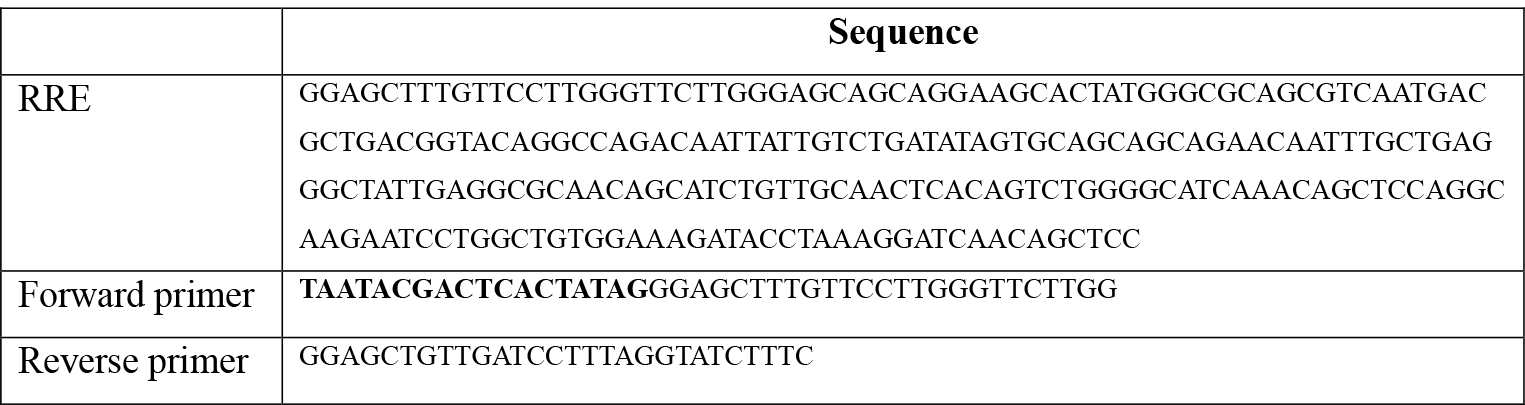

Both primers have an average melting temperature of 61°C but can vary. The forward primer contains a T7-promoter sequence that will be integrated into the amplified RRE fragment.

The PCR reagents used come from the Takara Bio Advantage^®^ 2 PCR kit and were mixed as follows:

- 8μl of DNA template (∼40ng)
- .5μl of both forward and reverse primer (10μM each)
- .5μl of 50X dNTP mix
- .5μl advantage PCR buffer
- .5μl 50X polymerase mix
- Complete to 25μl with nuclease-free water

Thermocycler program:

- 95°C - 1min
  - 95°C - 30sec
  - 68°C - 1min
- 68°C - 1min
- 4°C - hold

**Note**: The thermocycler protocol will vary depending on the specific PCR kit or reagents being used.

The PCR reaction is validated using a 2% agarose e-gel produced by Invitrogen. An optional step is to clean up the DNA using a ZYMO DNA clean and concentrator -5 kit and elute in 10μl of nuclease free water. This way the resulting DNA can be quantified, and specific amounts can be used for In-vitro transcription.

### In-vitro Transcription

NEB’s HiScribe T7 and Invitrogen’s MEGAscript are the two kits used for in-vitro transcription. Both follow the same protocol and there were no noticeable differences in yields.

The following reagents were mixed as follows:

- 2μl of ATP solution
- 2μl of CTP solution
- 2μl of GTP solution
- 2μl of UTP solution
- 2μl of 10X buffer
- 2ul of T7 enzyme mix
- 500ng – 1μg of DNA (up to 8μl)
- Complete to 20μl of nuclease-free water

**Note:** The reaction can be incubated anywhere between 3hrs to 16hrs at 37°C. For small fragments the typical incubation period should be longer. For the RRE a 16hr incubation was done, this can be followed by a 4°C hold overnight.

Afterwards, 1μl of TURBO DNase was added to the reaction and incubated at 37°C for 15min. The reaction was cleaned up using a ZYMO RNA clean and concentrator -25 kit.

**Tip**: For higher yields a -96 or -100 RNA clean and concentrator kit can be used. Additionally, when inserting the T7-promoter to the forward primer hav ing a run of two or three Gs increases transcription efficiency, so an additional G can be added to end of the promoter sequence: TAATACGACTCACTATAG(**G**)

The in-vitro transcription product can be validated using a 1% agarose Invitrogen e-gel or a 0.8% agarose gel. Concentrations are determined using the Thermo Scientific™ NanoDrop™ OneC Microvolume UV-Vis Spectrophotometer.

### DMS treatment

Make enough fresh refolding buffer (RB) for each of your samples (90-80μl per sample) before starting DMS and keep at 37°C. The following is an RB mix that can be upscaled or downscaled:

- 495 μL sodium cacodylate (0.4M)
- 3 μL MgCl_2_ (1M)

Prepare an aliquot of RNA that is between 1μg to 5μg in 10μl of nuclease-free water. Place the sample in 95°C for 1min. This step is essential for denaturing any secondary structures formed. Move the sample to 1.5ml tubes and add the RB and incubate for 20-30mins at 37°C.

**Note:** The amount of RB added will depend on the % of DMS used, which depends on the size of the RNA. For most cases, 1%-2% DMS is sufficient. Given that the final reaction volume is 100μl, 1% of DMS is equivalent to 1μl. This means that the volume of RB used is 90μl minus the % of DMS (**RB volume = 90μl – DMS%)**.

These next steps must be performed inside a chemical hood. Add x % DMS directly into the sample and incubate for 5 min at 37°C, shaking at 500RPM. For an untreated control sample add no DMS. Immediately add and mix 60 μl of Beta-mercaptoethanol (BME) directly into the solution to quench the reaction. The total volume (160μl) is used cleaned up using a ZYMO RNA clean and concentrator -5 kit. Alternatively, ethanol precipitation can be done by adding 3μl glycoblue, 18μl 3M Sodium Acetate, and 700μl 100% ice cold EtOH. This incubated at -80°C or in dry ice for 1hr or overnight. Spin down at max speed for 45 min at 4°C. Wash with 700 μl ice cold 75% EtOH and dry for 2-3mins. Resuspend in 10 μl of nuclease free water. Quantification of final DMS-modified RNA was done with the Thermo Scientific™ NanoDrop™ OneC Microvolume UV-Vis Spectrophotometer.

**Tip:** The amount of RNA can vary by molecule; some will require a higher input of RNA depending on how much is lost after DMS-treatment. It is possible to lose up to half of your starting input. This is due mainly to how the BME affects the buffers used for the ZYMO kit. For this protocol we recommend starting with 2μg of RNA, but this amount can be adjusted as long as the final concentration is ≥100ng/μl.

### Reverse transcription and cDNA PCR

Given that the RRE is smaller than 300bp, a simple RT-PCR protocol followed by NEBNext® Ultra™ II DNA Library Prep Kit can be used. For the RT-PCR the following reagents were mixed:

- ≥100ng of DMS-modified RNA
- 2μl of reverse primer
- 1μl of dNTPs
- .2μl of RNAseOUT^™^ (can substitute for other RNAse inhibitors)
- 4μl of NEB Induro^®^ reaction buffer
- 1μl of NEB Induro^®^ Reverse transcriptase
- Complete up to 20μl with nuclease free water

This reaction is incubated at 55°C for 10mins and then 95°C for 1min. Afterwards 1μl of 4M NaOH is mixed in and incubated at 95°C for 3mins to degrade any leftover RNA. An optional step is to clean up the RT-product using a ZYMO oligo clean and concentrator kit and eluted in 10μl of nuclease free water. The resulting cDNA is amplified by using the Takara Bio Advantage^®^ 2 PCR kit:

- 2μl of RT-product
- .5μl of both forward and reverse primer (10μM each)
- 2.5μl of 50X dNTP mix
- 2.5μl advantage PCR buffer
- .5μl 50X polymerase mix
- Complete to 26μl with nuclease-free water

Thermocycler program:

- 95°C - 1min
  - 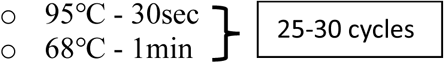
- 68°C - 1min
- 4°C - hold

The resulting cDNA was cleaned up using a ZYMO oligo clean and concentrator kit and eluted in 10μl of nuclease free water. A 2% agarose e-gel was used to validate the product and quantified as previously stated.

**Tip:** It is possible for the RT-PCR to have low or no yields. This is due to the Reverse transcriptase dropping off when there are two or more methylated bases adjacent to one another. In these cases, we recommend reducing the DMS% or increasing the input into the RT-PCR.

As stated earlier, the NEBNext® Ultra™ II DNA Library Prep Kit and its protocol were used to prepare the cDNA for sequencing. ∼500ng of cDNA were used as the starting material for library prep but amounts can vary. The only modification made to the protocol was skipping step 3 entirely, which consisted of size selection.

The final product was quantified using the Qubit4 fluorometer using Invitrogen’s 1x dsDNA HF kit. It is recommended to use this method of quantification instead of Nanodrop due to its precision.

### Sequencing and data processing

Samples were paired-end sequenced 150x150nt using the Illumina iSeq and NextSeq 1000. The sequencing quality and DMS-reactivities were analyzed using the DREEM algorithm (https://github.com/rouskinlab/DREEM) as in previous work^23^. The RNA 2D-structure illustrations were designed using a combination of VARNA, RNArtist (https://github.com/fjossinet/RNArtist) and Adobe Illustrator. Scatterplots and histogram were designed with Excell.

## Method II: DMS-MaPseq on PANDA with ASOs

### PCR and in-vitro transcription of PANDA

PANDA was PCR amplified from human Genomic DNA commercially available through Promega. Primers and ASOs were designed using SnapGene and ordered through IDT:

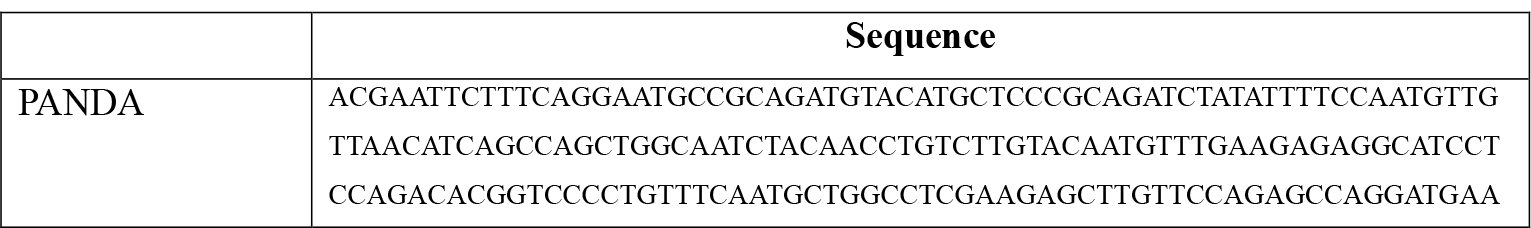
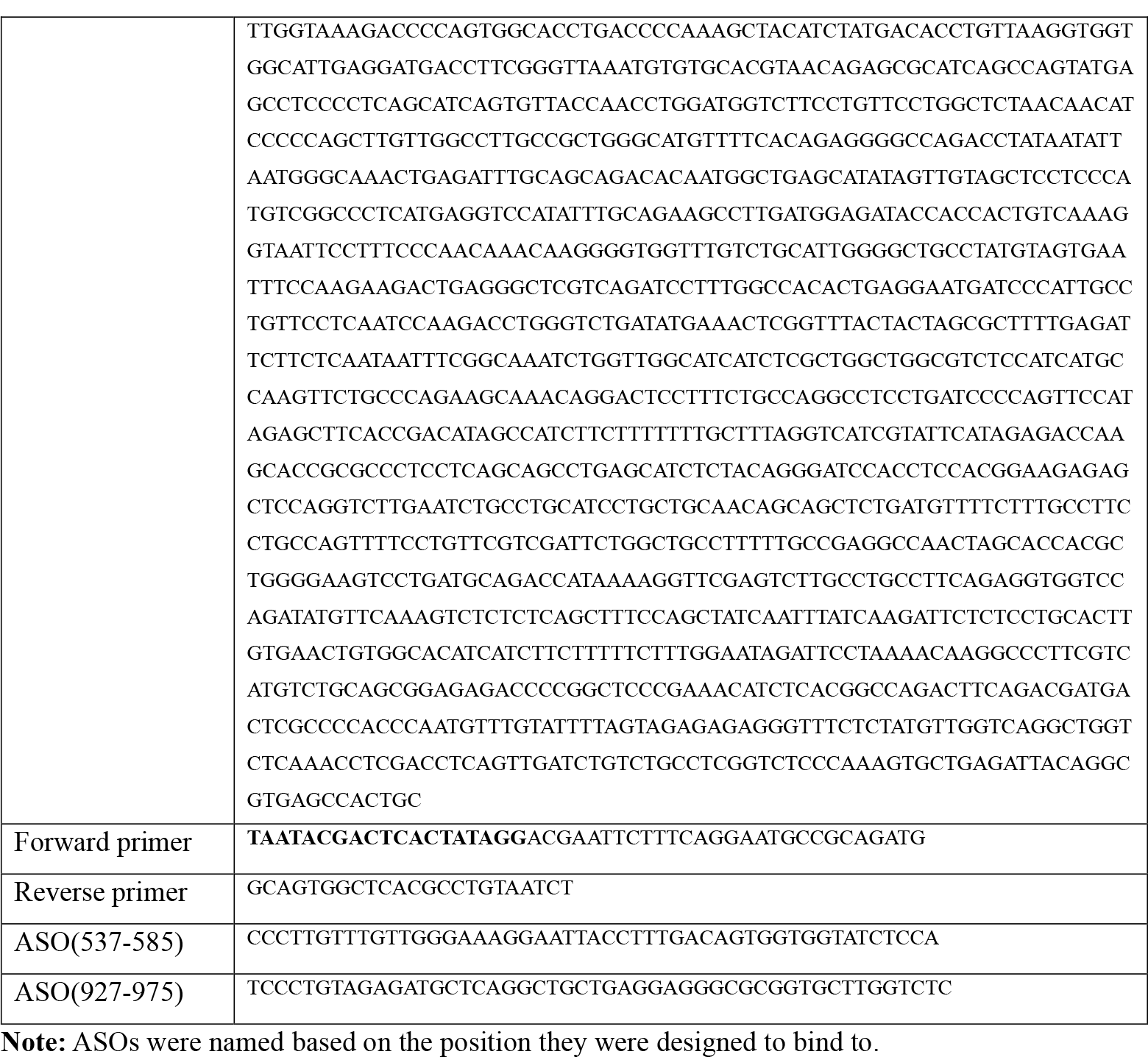

Both primers have an average melting temperature of 63°C but can vary. The forward primer contains a T7-promoter sequence that will be integrated into the PANDA PCR product.

A PCR master mix commercially available from Syd Labs, Inc was used. The reagents were mixed as follows:

- 3μl of genomic DNA
- 1μl of both forward and reverse primer (10μM each)
- 10μl PCR master mix
- Complete to 20μl with nuclease-free water

Thermocycler program:

- 95°C - 3min
  - 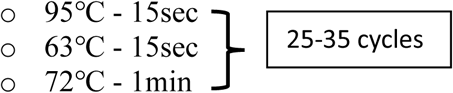
- 72°C - 5min
- 4°C - hold

**Note**: The thermocycler protocol will vary depending on the specific PCR kit or reagents being used.

The PCR reaction is validated using a 2% agarose e-gel produced by Invitrogen. An optional step is to clean up the DNA using a ZYMO DNA clean and concentrator -5 kit and elute in 10μl of nuclease free water. This way the resulting DNA can be quantified, and specific amounts can be used for In-vitro transcription. The in-vitro transcription protocol is the same as the one mentioned previously for the RRE, only difference is that it can be incubated for 3-4hr and still get relatively high yields.

### DMS and ASO treatment

Make enough fresh refolding buffer (RB) for each of your samples (90-80μl per sample) before starting DMS and keep at 37°C. The following is an RB mix that can be upscaled or downscaled:

- 495 μL sodium cacodylate (0.4M)
- 3 μL MgCl_2_ (1M)

Prepare you RNA and ASO sample by adding the following components into a 1.5ml tube:

- 10μl of (5 uM) ASO
- 1-5 pmol of your given RNA molecule
- Complete to 13.5μl with low EDTA TE buffer

The amount of ASO or RNA can be adjusted depending on the purpose and specificity of the experiment. For this protocol as little as 10X excess of ASO to RNA was sufficient to detect ASO binding. Place the sample in 95°C for 1min. Move the sample to 1.5ml tubes and add the RB and incubate for 20-30mins at 37°C.

**Note:** The sample volume for this protocol is different from the standard in-vitro DMS treatment, thus the volume of RB used will be different. Using the same approach as discussed previously the volume of RB used is 86.5μl minus the % of DMS (**RB volume = 86**.**5μl – DMS%)**.

These next steps must be performed inside a chemical hood. Add x % DMS directly into the sample and incubate for 5 min at 37°C, shaking at 500RPM. For an untreated control sample add no DMS. Immediately add and mix 60 μl of Beta-mercaptoethanol (BME) directly into the solution to quench the reaction. The total volume (160μl) is used cleaned up using a ZYMO RNA clean and concentrator -5 kit and resuspended in 20 μl of nuclease free water. Quantification of final DMS-modified RNA was done with the Thermo Scientific™ NanoDrop™ OneC Microvolume UV-Vis Spectrophotometer. This measurement will also pick up the concentration of ASO in your sample.

Afterwards, samples must undergo ASO degradation by mixing the following components in a 1.5ml tube and incubating at 37°C for 30mins:

- DMS-modified nucleic acid (∼2,000ng)
- 5μl of 10X TURBO DNase buffer
- 1μl of TURBO DNase enzyme
- Complete to 50μl of nuclease-free water

Samples can then be cleaned up using a ZYMO RNA clean and concentrator -5 kit and resuspended in 10 μl of nuclease free water.

### Sequencing library preparation

The library preparation kit used was xGen Broad-Range RNA Library Prep Kit distributed by IDT. It’s important to note that all volumes for each master mix is for one sample, the amounts need to be adjusted to the number of samples.

#### Fragmentation and Reverse Transcription

Prepare >100ng of DMS-modified RNA in 8μl in nuclease-free water. Preheat the thermocycler to 94°C and prepare the fragmentation mix by adding:

- 8μl RNA
- 1μl F1 (Random Primers)
- 4μl F3
- 2μL F4

Incubate fragmentation mix at 94ºC for 2mins and then place on ice immediately. Prepare the RT Mix without dNTPs by mixing:

- 2μl Induro RT
- 1μl R1 (RNAse Inhibitor)
- 1μl DTT

Add RT Mix to fragmentation reaction and incubate at room temperature for 30 minutes. This will allow the Induro RT to bind to the RNA/primer duplex before reverse transcription can start. After the 30 minutes add 2μL of F2 (dNTPs) and program the thermocycler as follows:

- 20ºC (hold to pre-heat)
- 20ºC for 10 min
- 42°C for 10min
- 55ºC for 60min
- PAUSE
  - Add 1μl 4M NaOH
- 95ºC for 3min
- 4ºC hold

Add 1μL of 4M HCl and complete the final volume to 50μL with low EDTA TE.

**Note:** Samples can be left in the 4°C overnight before adding the HCl

#### Ampure XP Bead Cleanup

Ampure XP beads can be exchanged for SPRI beads if needed.

Mix 50μL of Ampure XP Beads to samples and incubate samples at Room Temp for 5 minutes. Place samples on a magnetic rack for ∼2mins or until the solution has cleared. Remove the supernatant still on the magnetic rack without disturbing the brown pellet. Gently wash the pellet by adding 200μL of 80% Ethanol. Incubate for 30secs and pipette the EtOH out. Repeat the 80% Ethanol Wash. Make sure to remove as much EtOH as possible and incubate the beads on the magnet for 1min at room temperature. Once all the ethanol has dried, resuspend in 12μL of low EDTA TE and incubate for 2 minutes off the magnet. Place the tubes on the magnet again and incubate for ∼2 minutes until the solution is clear. Transfer 10μL to a new 0.2mL PCR tube. This is a stopping point and samples can be stored at -20ºC.

**Note:** When doing washes makes sure not to disrupt the pellet, otherwise most if not all of your library will be lost.

#### Adaptase Reaction

Prepare the adaptase Master Mix on ice by mixing the following reagents:

- 2μL Buffer A1
- 2μL Reagent A2
- 1.25μL Reagent A3
- 0.5μL Enzyme A4
- 0.5μL Enzyme A5
- 4.25μL Low-EDTA TE

For a final volume of 10.5μl per sample.

Before adding the adaptase master Mix, incubate samples at 95ºC for 2 min and transfer to ice immediately. add 10.5μL of master mix to each of your samples and mix thoroughly.

Run the following thermocycler program (Lid heat on):

- 37ºC for 15 min
- 95°C for 2min
- 4ºC hold

#### Extension Reaction

Prepare the extension master mix on ice by mixing the following:

- 1μL Reagent E1
- 22μL PCR Master Mix

For a final volume of 23μL per sample.

Add the extension master mix to samples and run the following thermocycler program (Lid Heat On):

- 98ºC for 1min
- 63ºC for 2 min
- 72ºC for 5 min
- 4ºC hold

#### Post-Extension Cleanups

Mix 52.5μL of SPRI Beads to samples and incubate samples at Room Temp for 5 minutes. Place samples on a magnetic rack for ∼2mins or until the solution has cleared. Remove the supernatant still on the magnetic rack without disturbing the brown pellet. Gently wash the pellet by adding 200μL of 80% Ethanol. Incubate for 30secs and pipette the EtOH out. Repeat the 80% Ethanol Wash. Make sure to remove as much EtOH as possible and incubate the beads on the magnet for 1min at room temperature. Once all the ethanol has dried, resuspend in 52μL of low EDTA TE and incubate for 2 minutes off the magnet. Place the tubes on the magnet again and incubate for ∼2 minutes until the solution is clear. Transfer 50μL to a new 0.2mL PCR tube.

#### Second SPRI Cleanup

Mix 60μL of SPRI Beads to samples and incubate samples at Room Temp for 5 minutes. Place samples on a magnetic rack for ∼2mins or until the solution has cleared. Remove the supernatant still on the magnetic rack without disturbing the brown pellet. Gently wash the pellet by adding 200μL of 80% Ethanol. Incubate for 30secs and pipette the EtOH out. Repeat the 80% Ethanol Wash. Make sure to remove as much EtOH as possible and incubate the beads on the magnet for 1min at room temperature. Once all the ethanol has dried, resuspend in 17μL of low EDTA TE and incubate for 2 minutes off the magnet. Place the tubes on the magnet again and incubate for ∼2 minutes until the solution is clear. Transfer 15μL to a new 0.2mL PCR tube. This is a stopping point and samples can be stored at -20ºC.

#### Ligation Reaction

Prepare the ligation master mix on ice by mixing the following:

- 3μL Buffer L1
- 10μL Reagent L2
- 2μL Enzyme L3

Mix the 15μl of ligation master mix with the sample and run the following thermocycler program (Lid Heat On):

- 25ºC for 15min
- 4ºC hold

#### Post-ligation Cleanup

Mix 30μL of SPRI Beads to samples and incubate samples at Room Temp for 5 minutes. Place samples on a magnetic rack for ∼2mins or until the solution has cleared. Remove the supernatant still on the magnetic rack without disturbing the brown pellet. Gently wash the pellet by adding 200μL of 80% Ethanol. Incubate for 30secs and pipette the EtOH out. Repeat the 80% Ethanol Wash. Make sure to remove as much EtOH as possible and incubate the beads on the magnet for 1min at room temperature. Once all the ethanol has dried, resuspend in 22μL of low EDTA TE and incubate for 2 minutes off the magnet. Place the tubes on the magnet again and incubate for ∼2 minutes until the solution is clear. Transfer 20μL to a new 0.2mL PCR tube. This is a stopping point and samples can be stored at -20ºC.

#### Indexing PCR

Prepare the indexing PCR master mix on ice by mixing the following:

- 25μL PCR Master Mix
- 5μL single index primer (dual index primers can also be used)

Mix master mix with your sample and run the following thermocycler program:

- 98ºC for 2min
  - 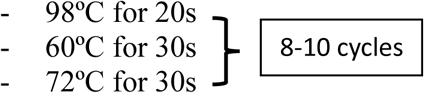
- 72ºC for 1min
- 4ºC hold

This is a stopping point and samples can be stored at -20ºC.

**Hint**: If yields after library preparation are too low you can try increasing the number of cycles for the indexing PCR up to 12.

#### Library Size Selection and Gel Extraction

Prepare 20μL of the indexing PCR reaction by adding 4μL of 6x Loading Dye. Load the entirety into an 8% TBE Gel, which are commercially available through Invitrogen. Load 4μL of 1kb plus ladder and run for 55m @ 180V with 1X TAE running buffer. Afterwards place the gel in TBE Buffer with SYBR Gold and incubate for ∼5mins. Gently place the gel on a UV light box or a blue light transilluminator and excise the libraries that are between the 250bp and 600bp marker. Adapters will be present around 200bp so those should be avoided. Place the gel fragments into punctured 0.5mL tubes placed inside 1.5mL tube and spin down for 1min at maximum speed (∼21,000g) to crush the gel through the hole. Discard the punctured tube and add 500μL 300mM NaCl to the crushed gel. Incubate at 70ºC 1500RPM for 20mins. Pipette total gel slurry into a costar column and centrifuge at max speed for 1min. Move the eluent to a new tube and add 600μL of isopropanol + 3μL of glycoblue. Incubate samples at -20ºC for at least 1h but they can be left overnight. Centrifuge at max speed (∼18,000g) for 45min at 4ºC. Remove the supernatant without disrupting the pellet and gently wash with 500μL ice-cold 70% EtOH. Aspirate all the EtOH without disrupting the pellet and leave the tubes open at room temperature for 1-2mins for the residual EtOH to evaporate. Resuspend the pellet in 15μL nuclease-free water. The final product was quantified using the Qubit4 fluorometer using Invitrogen’s 1x dsDNA HF kit. It is recommended to use this method of quantification instead of Nanodrop due to its precision. It is expected for samples to have a concentration of 5-15ng/μl but having a concentration as low as 1.0ng/μl will be enough for sequencing.

### Sequencing and data processing

Samples were paired-end sequenced 150x150nt using the Illumina iSeq and NextSeq 1000. The sequencing quality and DMS-reactivities were analyzed using the DREEM algorithm (https://github.com/rouskinlab/DREEM) as in previous work^23^. The RNA 2D-structure illustrations were designed using a combination of VARNA, RNArtist (https://github.com/fjossinet/RNArtist), and Adobe Illustrator. Scatterplots were designed with Excell. AUROC curve and R^2^ sliding windows were designed with custom Python scripts.

## Results

### HIV1-RRE alternative structure

A bar graph for the mutational fraction was generated for three separate samples, two 1% DMS and a 1.5% DMS treatment (Fig 1A). All three samples correlate well, demonstrating the robustness and reproducibility of our method (Fig 1B). After running DREEM on the samples, the number of mutated bases per read count was plotted (Fig 1C). For one of our 1% DMS duplicates we see that most reads have 2-3 mutations, an ideal profile for clustering. These samples were sequenced deeper to generate the clustered profiles, predicting two alternative structures as expected (Fig 1D). Like in previous work, we demonstrate that the HIV1-RRE adopts a five-stem and a four-stem structure^28^ (Fig 1E). These results confidently show that the updated DMS-MaPseq protocol works as expected for in-vitro RNA structure prediction.

**Figure 1:**
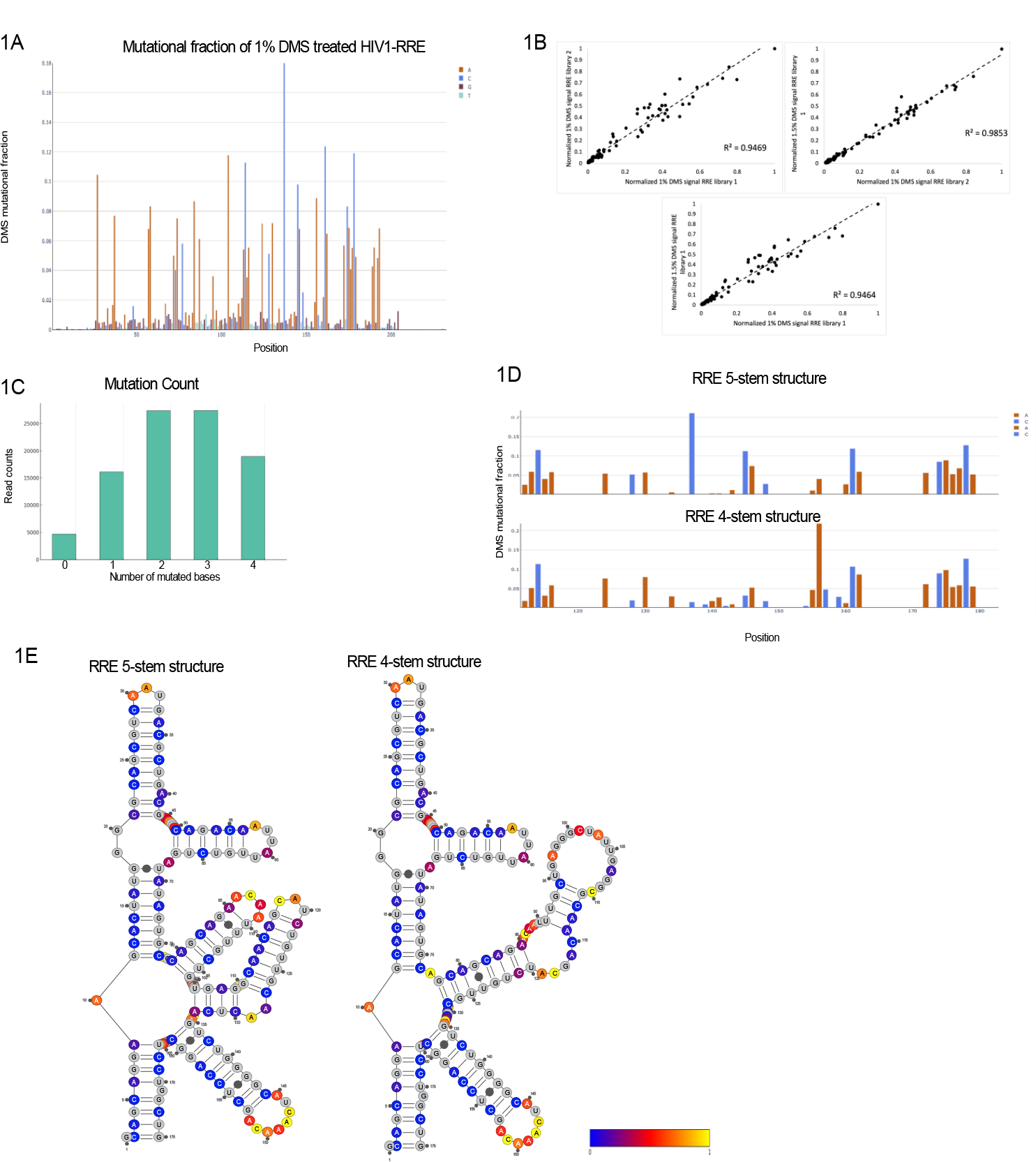
DMS-MaPseq generated data and predicted structure of the HIV1-RRE. **A**) Bar graph of the mutational fraction for the 1% DMS treated RRE. Unpaired adenines (red) and cytosines (blue) have a high mutational fraction, as opposed to guanines (brown), uridines (cyan) or paired bases. **B**) Scatter plot of the DMS signal for each replicate. Normalized values represent mutated A or Cs after masking Us and Gs. **C**) Histogram of mutations per read count of a 1% DMS treated sample. **D**) Bar graphs of the mutational fractions generated for both predicted clusters. Unpaired adenines (red) and cytosines (blue) have a high mutational fraction, as opposed to guanines (brown), uridines (cyan) or paired bases. **E**) 2D structure predictions of the 5-stem RRE (left), 4-stem RRE (right). Bases are color-coded by DMS reactivity; yellow being highest reactivity, blue lowest, and gray indicating masked positions.

### DMS-MaPseq resolved structure of PANDA

The structure of PANDA lncRNA was generated using DMS reactivities as constraints in RNAstructure^29^ (Fig 2A). The normalized DMS reactivity for two replicates were plotted and yielded an R^2^ of 0.99, indicating high correlation between both samples (Fig 2B). We used an AUROC curve to quantify how well model fits the data (Fig 2C). The calculated AUC is 0.91, indicating that the DMS reactivities highly agree with the structure model.

**Figure 2:**
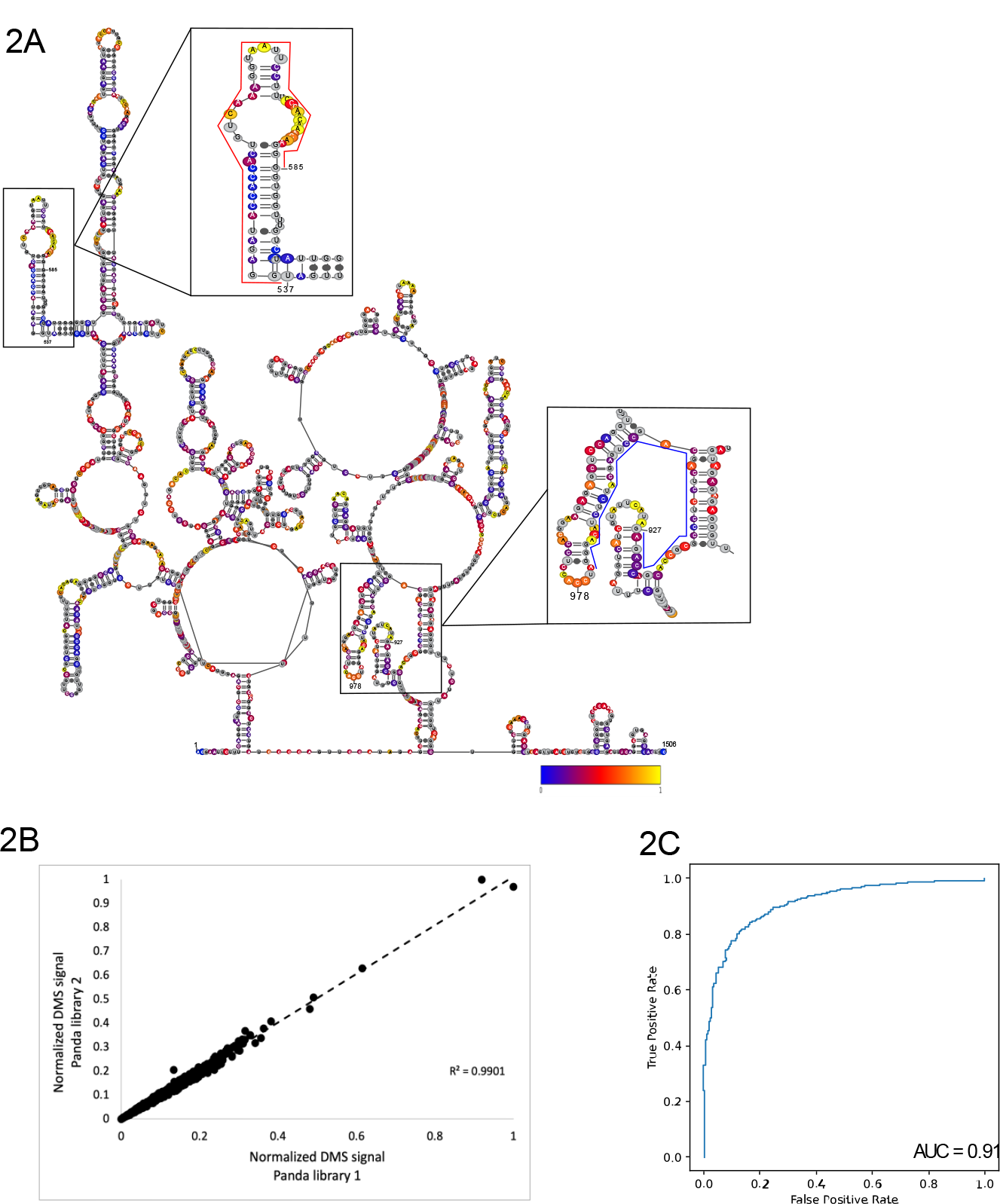
DMS-MaPseq generated data and predicted structure of the PANDA lncRNA. **A**) Predicted 2D structure ensemble for the average DMS reactivities of in-vitro folded PANDA. Target of ASO(537-585) is encased in red. Target of ASO(927-975) is encased in blue. Bases are color-coded by DMS reactivity; yellow being highest reactivity, blue lowest, and gray indicating masked positions. **B**) Scatter plot of the DMS signal for each replicate. Normalized values represent mutated A or Cs after masking Us and Gs. **C**) AUROC and AUC values generated from the structure ensemble data and normalized DMS reactivities after masking Gs and Us.

### Detection of ASO binding to PANDA

The normalized DMS reactivities between replicates treated with each ASO respectively corelate well as evident by an R^2^ of 0.996 for samples treated with ASO(537-585) and 0.976 for samples treated with ASO(927-975) (Fig 3A). ASO binding was detected as a loss in DMS reactivities of the bases with which the ASO is base pairing with. This can be seen for samples treated with either of the two ASOs (Fig 3B, 3D). A sliding window that compares the R^2^ between the untreated and ASO treated samples was used to measure the change in DMS signal. For samples treated with ASO(537-585) there is an extreme drop in the R^2^ at positions 500-600, which agrees well with the drop in DMS reactivity at the ASO binding site (Fig 3C). Unexpectedly, there is also a slight decrease around positions 800. This could be due either to an off-target effect, or perhaps there is a long-distance interaction occurring between both regions of the RNA which is disrupted by the ASO. Samples treated with ASO(927-975) show a clear drop in the R^2^ at positions 900-1,000 just as expected (Fig 3E). These experiments demonstrated how DMS-MaPseq can be used to detect binding of ASOs to specific sites of a given RNA.

**Figure 3:**
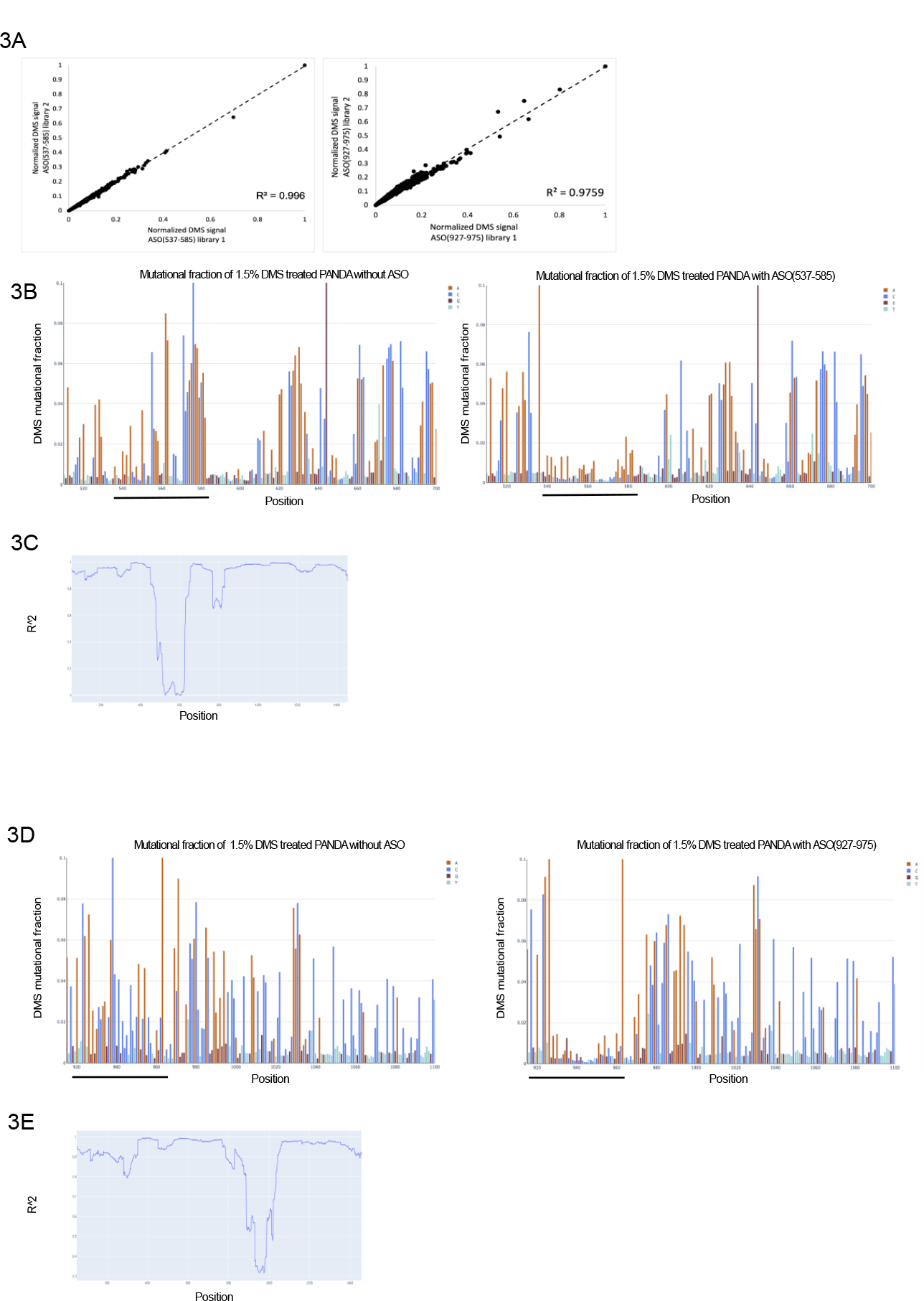
In-vitro DMS-MaPseq detection of ASO binding to PANDA. **A**) Scatter plot for the DMS signal for replicates treated with ASO(537-585) (left) or ASO(927-975) (right). Normalized values represent mutated A or Cs after masking Us and Gs. **B**) Bar graph of the mutational fraction for the 1.5% DMS treated Panda without ASO (left) and with ASO(537-585) (right). Unpaired adenines (red) and cytosines (blue) have a high mutational fraction, as opposed to guanines (brown), uridines (cyan) or paired bases. The ASO binding sight is indicated by the black line under the graph. **C**) Sliding window comparing the R^2^ values of the DMS reactivities between PANDA without and with ASO(537-585) treatment. A 100-nucleotide window was used. **D**) Bar graph of the mutational fraction for the 1.5% DMS treated Panda without ASO (left) and with ASO(927-575) (right). Unpaired adenines (red) and cytosines (blue) have a high mutational fraction, as opposed to guanines (brown), uridines (cyan) or paired bases. The ASO binding sight is indicated by the black line under the graph. **E**) Sliding window comparing the R^2^ values of the DMS reactivities between PANDA without and with ASO(537-585) treatment. A 100-nucleotide window was used.

## References

1. Mortimer, S.A., Kidwell, M.A. & Doudna, J.A. Insights into RNA structure and function from genome-wide studies. Nat. Rev. Genet. 15, 469–479 (2014).

2. Leppek, K., Das, R. & Barna, M. Functional 5′ UTR mRNA structures in eukaryotic translation regulation and how to find them. Nat. Rev. Mol. Cell Biol. 19, 158–174 (2018).

3. Mayr, C. Regulation by 3′-untranslated regions. Annu. Rev. Genet. 51, 171–194 (2017).

4. Spitale, R.C., Incarnato, D. Probing the dynamic RNA structurome and its functions. Nat Rev Genet 24, 178–196 (2023).

5. RNA levers and switches controlling viral gene expression.

6. Ma, H., Jia, X., Zhang, K. et al. Cryo-EM advances in RNA structure determination.

7. Fallmann J, Will S, Engelhardt J, Gruning B, Backofen R, Stadler PF, Recent advances in RNA folding, J Biotechnol 261 (2017) 97–104. [PubMed: 28690134]

8. Ehresmann C, Baudin F, Mougel M, Romby P, Ebel JP, Ehresmann B, Probing the structure of RNAs in solution, Nucleic Acids Res 15(22) (1987) 9109–28. [PubMed: 2446263]

9. Wells, S.E., Hughes, J.M., Igel, A.H. & Ares, M. Jr. Use of dimethyl sulfate to probe RNA structure in vivo. Methods Enzymol. 318, 479–493 (2000).

10. Zubradt, M., Gupta, P., Persad, S. et al. DMS-MaPseq for genome-wide or targeted RNA structure probing in vivo. Nat Methods 14, 75–82 (2017).

11. Inoue T, Cech TR, Secondary structure of the circular form of the Tetrahymena rRNA intervening sequence: a technique for RNA structure analysis using chemical probes and reverse transcriptase, Proc Natl Acad Sci U S A 82(3) (1985) 648–52. [PubMed: 2579378]

12. Tijerina P, Mohr S, Russell R, DMS footprinting of structured RNAs and RNA-protein complexes, Nat Protoc 2(10) (2007) 2608–23. [PubMed: 17948004].

13. Homan, P. J. et al. Single-molecule correlated chemical probing of RNA. Proc. Natl Acad. Sci. USA 111, 13858–13863 (2014).

14. Mohr S, Ghanem E, Smith W, Sheeter D, Qin Y, King O, Polioudakis D, Iyer VR, Hunicke-Smith S, Swamy S, Kuersten S, Lambowitz AM. Thermostable group II intron reverse transcriptase fusion proteins and their use in cDNA synthesis and next-generation RNA sequencing. RNA. (2013)

15. Strobel, E. J., Yu, A. M. & Lucks, J. B. High-throughput determination of RNA structures. Nat. Rev. Genet. 19, 615–634 (2018).

16. Lan, T.C.T., Allan, M.F., Malsick, L.E. et al. Secondary structural ensembles of the SARS-CoV-2 RNA genome in infected cells. Nat Commun 13, 1128 (2022).

17. Tomezsko, P.J., Corbin, V.D.A., Gupta, P. et al. Determination of RNA structural diversity and its role in HIV-1 RNA splicing. Nature 582, 438–442 (2020).

18. Morandi, E., Manfredonia, I., Simon, L.M. et al. Genome-scale deconvolution of RNA structure ensembles. Nat Methods 18, 249–252 (2021).

19. Olson SW, Turner AW, Arney JW, Saleem I, Weidmann CA, Margolis DM, Weeks KM, Mustoe AM. Discovery of a large-scale, cell-state-responsive allosteric switch in the 7SK RNA using DANCE-MaP. Mol Cell. 2022 May 5;82(9):1708-1723.e10.

20. Crooke ST, Liang XH, Baker BF, Crooke RM. Antisense technology: A review. J Biol Chem. 2021 Jan-Jun; 296:100416.

21. Dhuri K, Bechtold C, Quijano E, Pham H, Gupta A, Vikram A, Bahal R. Antisense Oligonucleotides: An Emerging Area in Drug Discovery and Development. J Clin Med. 2020 Jun 26;9(6):2004.

22. Sophie F. Hill, Miriam H. Meisler; Antisense Oligonucleotide Therapy for Neurodevelopmental Disorders. Dev Neurosci 10 September 2021; 43 (3-4): 247–252.

23. Hung T, Wang Y, Lin MF, Koegel AK, Kotake Y, Grant GD, Horlings HM, Shah N, Umbricht C, Wang P, Kong B, Langerod A, Borresen-Dale AL, Kim SK, van de Vijver M, Sukumar S, Whitfield ML, Kellis M, Xiong Y, Wong DJ and Chang HY: Extensive and coordinated transcription of noncoding RNAs within cell-cycle promoters. Nat Genet 43: 621–629, 2011.

24. Wang KC, Chang HY. Molecular mechanisms of long noncoding RNAs. Mol Cell. 2011 Sep 16;43(6):904–14.

25. Huang HW, Xie H, Ma X, Zhao F, Gao Y. Upregulation of LncRNA PANDAR predicts poor prognosis and promotes cell proliferation in cervical cancer. Eur Rev Med Pharmacol Sci. 2017 Oct;21(20):4529–4535.

26. Liu J, Ben Q, Lu E, He X, Yang X, Ma J, Zhang W, Wang Z, Liu T, Zhang J, Wang H. Long noncoding RNA PANDAR blocks CDKN1A gene transcription by competitive interaction with p53 protein in gastric cancer. Cell Death Dis. 2018 Feb 7;9(2):168.

27. Peng C, Hu W, Weng X, Tong R, Cheng S, Ding C, Xiao H, Lv Z, Xie H, Zhou L, Wu J, Zheng S. Over Expression of Long Non-Coding RNA PANDA Promotes Hepatocellular Carcinoma by Inhibiting Senescence Associated Inflammatory Factor IL8. Sci Rep. 2017 Jun 23;7(1):4186.

28. Tomezsko P, Swaminathan H, Rouskin S. Viral RNA structure analysis using DMS-MaPseq. Methods. 2020 Nov 1;183:68–75.

29. Reuter, J.S., Mathews, D.H. RNAstructure: software for RNA secondary structure prediction and analysis. BMC Bioinformatics 11, 129 (2010).

